# The Design Principles of Biochemical Timers: Circuits That Discriminate Between Transient and Sustained Stimulation

**DOI:** 10.1101/100651

**Authors:** Jaline Gerardin, Wendell A. Lim

## Abstract

Many cellular responses for which timing is critical display temporal filtering – the ability to suppress response until stimulated for longer than a given minimal time. Temporal filtering can play a key role in filtering noise, choreographing the timing of events, and mediating the interpretation of dynamically encoded signals. To define the biochemical circuits capable of kinetic filtering, we comprehensively searched the space of three-node networks. We define a metric of “temporal ultrasensitivity”, a measure of the steepness of activation as a function of stimulus duration. We identified five classes of core network motifs capable of temporal filtering, each with different functional properties such as rejecting high frequency noise, committing to response (bistability), and distinguishing between long stimuli. Combinations of the two most robust motifs, double inhibition (DI) and positive feedback with AND logic (PF_AND_), underlie several natural timer circuits involved in processes such as cell cycle transitions, T cell activation, and departure from the pluripotent state. The biochemical network motifs described in this study form a basis for understanding the common ways in which cells make dynamic decisions.

## INTRODUCTION

Timing is critical in biological regulation. How do cells tell time and measure the duration of events? In many processes, cells display *temporal filtering* or *temporal thresholding* -- the ability to measure the duration of time that they experience a given input, and to only respond after a given threshold duration of stimulation (**Fig 1A**). Closely related behavior has also been referred to as kinetic proofreading (Hopfield, 1974) or persistence detection (Mangan and Alon, 2003). Temporal filtering is important for several types of physiological behaviors. Signaling networks downstream of receptors must filter noisy, transient environmental fluctuations to distinguish them from real, more sustained signals (Hopfield, 1974). Kinetic filters can absorb and dissipate these transient inputs. Measuring stimulation time also allows cells to trigger a response to an initial cue only after a specific delay, which can be critical for coordinating the relative timing of events, especially in complex, sequential processes such as the cell cycle or development. Finally, there is increasing appreciation that biological information can be encoded dynamically, e.g. in features such as input duration or frequency (Batchelor et al., 2011; Locke et al., 2011; Purvis and Lahav, 2013; Suel et al., 2007), and cellular circuits that can measure duration of stimulation undoubtedly play a key role in interpreting and decoding this kind of more complex temporal information.

**Figure 1.**
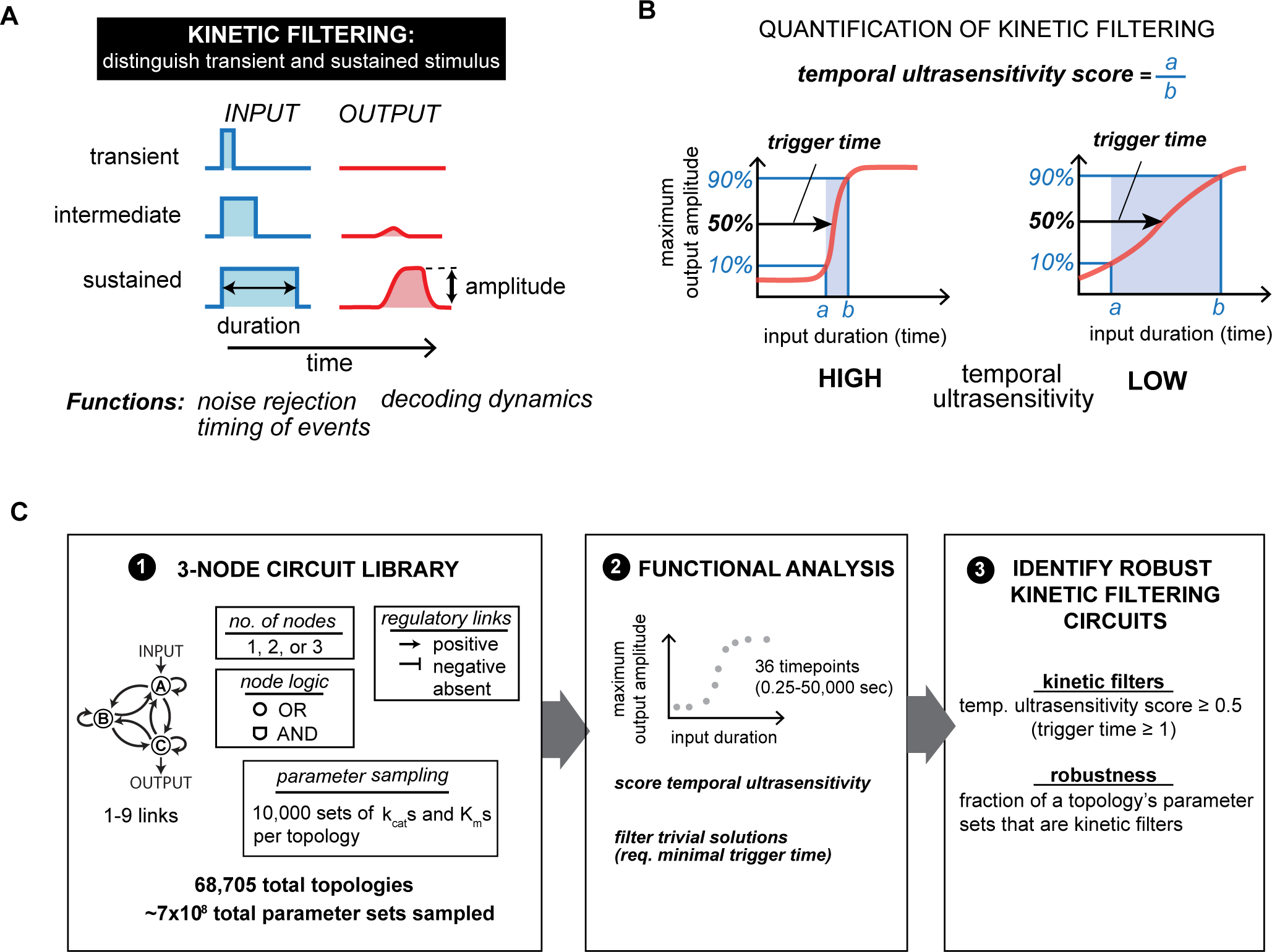
Kinetic filtering circuits distinguish between transient and sustained inputs. (A) Kinetic filtering circuits respond to sustained but not transient stimuli, allowing cells to perform time-sensitive functions. (B) Temporal ultrasensitivity score quantifies kinetic filtering by measuring steepness of activation over stimulus time, defined by taking the ratio of input duration required for 10% activation to input duration required for 90% activation. Trigger time, the duration of input yielding 50% activation, measures the duration of stimulus necessary to trigger response. (C) To identify kinetic filtering architectures, temporal ultrasensitivity score and trigger time were measured over an enumerated space of 68,705 circuit topologies and 10,000 sampled parameter sets per topology. Parameter sets were considered to show kinetic filtering if temporal ultrasensitivity score ≥ 0.5 and trigger time ≥ 1s. A topology’s robustness is defined as the fraction of its sampled parameter sets that show kinetic filtering. See Figure S1 for details on simulating enzymatic circuits with OR and AND nodes.

How can biochemical circuits function as kinetic filters? There have been few studies that systematically explore and compare which signaling circuit architectures can kinetically filter stimuli and measure time. Although some circuits that can serve as kinetic filters have been analyzed, for many biological examples, the precise molecular circuitry or mechanism responsible for the measurement of stimulus time is not known. Some specific classes of circuits have been noted to be able to serve as kinetic filters. These include extended multistep cascades (Hopfield, 1974; Samoilov et al., 2002), as well as coherent feed forward loops that have both long (multi-step) and short branches of transmission that are simultaneously required for output (Mangan and Alon, 2003; Murphy et al., 2002). In transcriptional networks, positive feedback has also been proposed to play a role in noise suppression (Horning and Barkai, 2008). But are these the only solutions for effective kinetic filtering? If there are more families of solutions, how do they compare with one another in terms of efficiency and various functional tradeoffs? As we begin to understand how the cell coordinates and interprets complex dynamic information, it will be important to have a road map to help identify and classify the general types of molecular circuits that will emerge.

Coarse-grained network enumeration offers a computational approach to identify classes of biochemical network architectures that can achieve a given target function (Chau et al., 2012; Lim et al., 2013; Ma et al., 2006; 2009). Comprehensive, unbiased enumeration of a space of simple circuits allows identification of core solutions and evolutionary starting points for more complex networks. A set of core solutions forms the basis for understanding and cataloging natural timing circuits as well as providing blueprints for design synthetic circuits that can measure time.

Here we apply this approach to search the full space of all possible 1-, 2-, and 3-node enzymatic networks and identify the classes of network architectures that can robustly achieve kinetic filtering. To identify kinetic filters, we defined a new metric – *temporal ultrasensitivity* – to measure the steepness with which activation of a system occurs as a function of increasing stimulus duration. As the name implies, temporal ultrasensitivity is an analog of concentration ultrasensitivity – the measure of the steepness of a system’s dose-response (Goldbeter and Koshland, 1981). Here we find five classes of simple network motifs that can achieve kinetic filtering, including the previously characterized coherent feed forward loop (Mangan and Alon, 2003; Mangan et al., 2003). Two of these motifs can be optimized to yield kinetic filters with both sharp temporal ultrasensitivity and long trigger time (duration of input required for half-maximal response). In contrast, the other motifs identified can robustly achieve high temporal ultrasensitivity only at a lower range of trigger times. We identify key mechanistic properties that allow for longer trigger times while retaining sharp activation dynamics.

These findings and the tradeoffs associated with each class of motif suggest particular functional roles of each subtype. Interestingly, one of the most common convergent motifs among natural kinetic filters, responsible for cell cycle transition circuits and T cell activation among other functions, is a combination of the two most robust classes uncovered in our search, which are predicted to combine both long, tunable trigger times with committed, temporally ultrasensitive switching. We also predict that other types of combined circuit motifs that would have particular kinetic filtering properties.

Our understanding of how cells respond to the complex dynamic information they receive, and how they control their own responses over time, is currently relatively primitive. The design principles of kinetic filtering circuits explored in this study may help provide an initial road map for uncovering, identifying and understanding such temporal regulatory mechanisms.

## RESULTS

### Defining kinetic filtering: temporal ultrasensitivity and trigger time

We use two parameters to quantitatively define a kinetic filter (**Fig 1B**). First, trigger time is the input duration required to achieve half-maximal response; second, temporal ultrasensitivity is the steepness of the response vs input duration curve. Analogous to concentration-based ultrasensitivity, temporal ultrasensitivity quantifies the sharpness of a signaling network’s kinetic filtering thresholding behavior, implying a steep temporal dose response curve where input stimulation with durations shorter than the trigger time result in minimal or no activation of the network, and inputs longer than the trigger time result in maximal activation. Networks that perform as kinetic filters can thus be defined as those that show temporal ultrasensitivity or trigger times above a minimum cutoff value.

### Circuit enumeration and analysis of robust kinetic filtering

To identify the simplest kinetic filtering circuits, we enumerated the space of 1-, 2-, and 3-node enzymatic circuits, as described previously (Ma et al., 2009), and measured each topology’s temporal ultrasensitivity and trigger time under multiple parameter sets (**Fig 1C**). Here we focused on enzymatic nodes, where each node is modeled with standard Michaelis-Menton parameters. Each node is at a fixed total concentration partitioned into active and inactive states. A regulatory link between nodes indicates that the active state of the upstream node catalyzes the conversion of the downstream node between its active and inactive state (**Fig S1**).

In this model, nodes that integrate two regulatory inputs exhibit “AND” or “OR” logic. Here we do not define these as absolute Boolean operators, but rather use this nomenclature to describe whether the effects of two different upstream regulatory links are either multiplicative (“AND”) or additive (“OR”) (**Fig S1**). Both logics are sampled in this enumeration because prior work has shown that some key kinetic filters require specific integrating node logic (Mangan and Alon, 2003). These classes of nodes only describe two extreme models of two-input integration where both inputs are absolutely needed or both equally activating, but many intermediate behaviors are also possible, such as integrating nodes in which the weights of activation by each link are different (one input link is weak activator; other is strong).

Altogether, we search a space of 68,705 possible network architectures and quantify temporal ultrasensitivity and trigger time for each architecture. We measure these behaviors by tracking maximal output at any time during the simulation, since we also wanted to be able to capture networks that may require a substantial delay after the stimulus pulse to develop its full output (note that kinetic filtering as defined here is focused only on the duration of input stimulation, and is agnostic about the time delay required to see output).

To capture only the kinetic filters with very strong all-or-none response to input duration, we applied a high stringency cutoff for temporal ultrasensitivity satisfied by <1% of simulated circuits (**Fig 2A**). Additionally, we required the circuits display a minimum trigger time of 1s (compared to tested input durations of up to 50,000 s) to exclude circuits with trivially short trigger times (**Fig S2A**). These cutoffs yield circuits with steep dynamic activation thresholds, but with a range of trigger times. We then quantified the *robustness* of each topology’s kinetic filtering by measuring the fraction of a topology’s tested parameter sets that satisfy the performance cutoffs (Dassow et al., 2000; Hornung and Barkai, 2008; Ma et al., 2009). A higher robustness implies that the topology’s performance is more robust to changes in parameter values. High robustness circuits are thought to represent the most likely solutions to emerge from a random evolutionary process (Lim et al., 2013).

**Figure 2.**
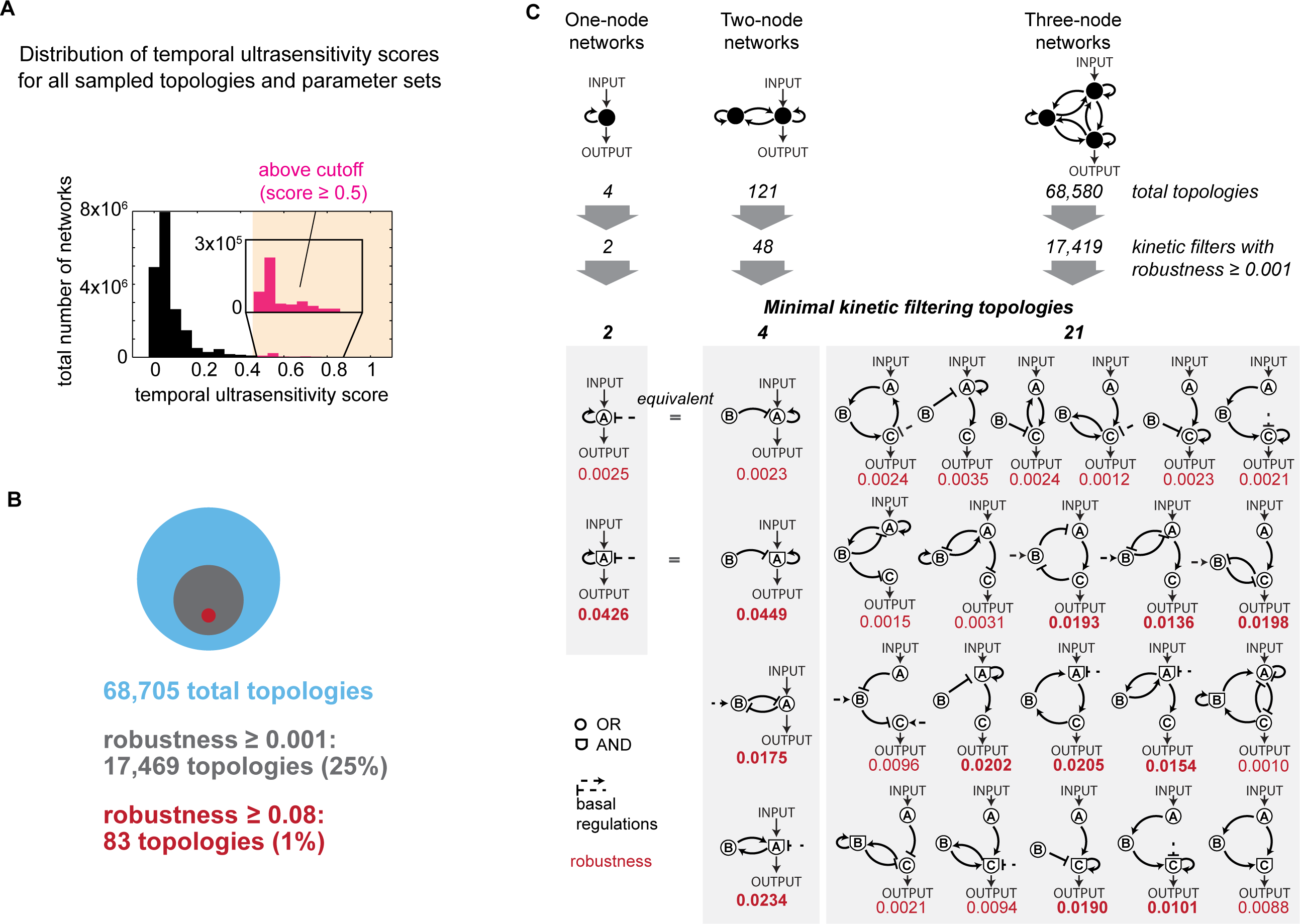
Enumeration of 1-, 2-, and 3-node networks finds 25 minimal kinetic filtering circuits. (A) Distribution of temporal ultrasensitivity scores across all topologies and parameter sets. A stringent cutoff of temporal ultrasensitivity score ≥ 0.5 identifies circuits capable of kinetic filtering. See Figure S2A for distribution of trigger times among circuits with temporal ultrasensitivity score ≥ 0.5. (B) Out of 68,705 total topologies, 25% have robustness above a low cutoff of 0.001(at least 10/10,000 parameter sets satisfy temporal ultrasensitivity score ≥ 0.5 and trigger time ≥ 1s) and 1% have robustness above a high cutoff of 0.08 (at least 800/10,000 parameter sets satisfy temporal ultrasensitivity score ≥ 0.5 and trigger time > 1s). (C) Number of topologies, kinetic filters with robustness ≥ 0.001, and minimal kinetic filters in 1-, 2-, and 3-node networks. Minimal kinetic filtering topologies are topologies with robustness ≥ 0.001 where removal of any link decreases robustness below 0.001 (Figure S2B). Two 2-node minimal kinetic filters are topologically identical to 1-node minimal kinetic filters with regulatory node B taking the place of the basal regulator.

We explored network space using a two-phase search strategy. First we searched the network space with a low-stringency robustness cutoff, where only very low-robustness networks were eliminated, in order to cast a wide net to find all minimal network motifs than can perform kinetic filtering. We can search this set of architectures for clusters of core kinetic filtering network motifs. In the second phase, we searched for optimal kinetic filters, which are likely to be more complex networks, by applying a much higher robustness cutoff. We could then identify whether particular kinetic filtering motifs identified in the first search were enriched within this more selective set of network architectures.

Over 25% of all the enumerated topologies satisfied the low stringency robustness cutoff, but only 83 topologies met the high robustness cutoff (**Fig 2B**). We first describe phenotypic clustering of the large set of topologies and define the main classes of minimal kinetic filtering motifs, then explore motif enrichment in the smaller, higher-performing set of topologies.

### Low-stringency search and phenotypic clustering identifies five classes of kinetic filtering networks

Of the 17,469 topologies that passed the low-stringency robustness cutoff, many are likely to be redundant – closely related networks that contain the same core motif that executes kinetic filtering. To reduce this large set of topologies to a minimal core set of non-redundant architectures, we systematically tested the effect of link pruning on robustness (Chau et al., 2012) (**Fig S2B**). We considered a minimal kinetic filtering architecture to be one in which removal of any single link in the network resulted in a drop in robustness below the 0.001 cutoff.

After this pruning procedure, 25 topologies were classified as minimal kinetic filtering architectures (**Fig 2C**). Each of the 17,469 kinetic filters in the low-stringency robustness set contains at least one of these 25 minimal kinetic filters as a core substructure, and the most robust kinetic filters often contain multiple minimal architectures. The minimal architectures thus form a basis set for analysis and classification of kinetic filters.

To investigate whether the minimal topologies could be clustered by functional behaviors, we measured six “phenotypic” metrics for each parameter set of all the minimal topologies that were above our kinetic filtering metric cutoffs, a total of 2896 topology/parameter combinations. The phenotypic metrics (**Fig 3A**) were: 1) trigger time, the duration of input required to achieve 50% maximal activation, without regard to kinetics of activation; 2) whether the network exhibited long-term memory, defined as final output concentration divided by maximum output concentration; 3) time for turning the circuit on (the time required for output to reach 50% maximum amplitude after application of input; 4) steepness of the circuit turning on (time required for output to reach 10% maximum amplitude divided by time required to reach 90% amplitude); 5) time to turn the circuit off (time required for output to decrease to 50% maximum amplitude after the input was turned off); and 6) steepness of the circuit turning off (time required for output to decrease to 10% maximum amplitude after the input was turned off divided by time required to reach 90% of maximum amplitude after the input was turned off). Metrics 2-6 were measured for a single input pulse of duration 50,000 seconds.

**Figure 3.**
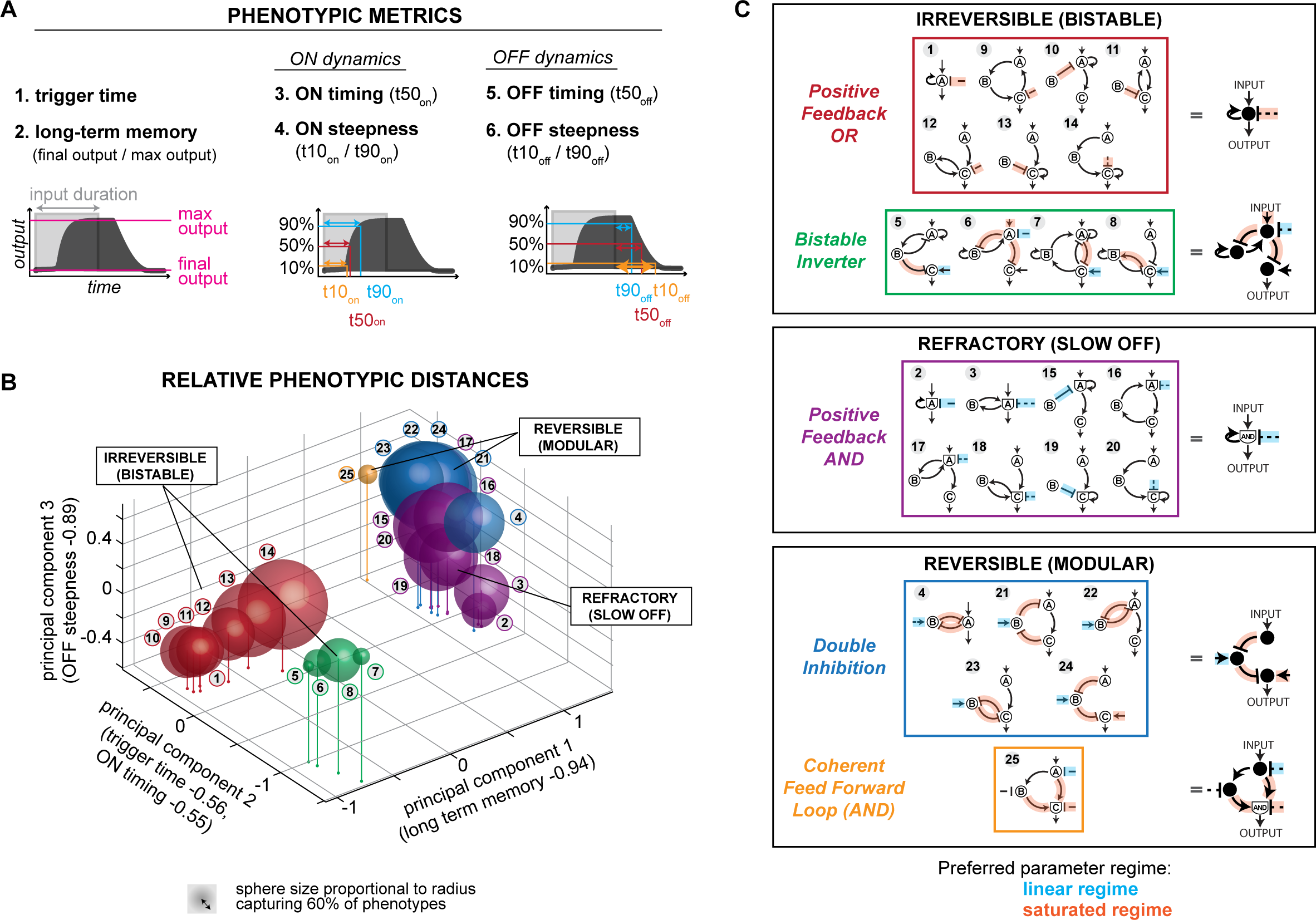
Minimal kinetic filters of 1, 2, and 3 nodes phenotypically cluster into five groups. (A) Metrics used to cluster minimal kinetic filters by phenotypic features. For metrics 2-5, a single input pulse of duration 50,000s was applied. Measurements of ON dynamics are relative to input ON time, and measurements of OFF dynamics are relative to input OFF time. OFF dynamics were not measured for circuits where max output = final output. Phenotypic metrics were measured for all parameter sets of minimal kinetic filters with temporal ultrasensitivity ≥ 0.5 and trigger time ≥ 1s (total of 2,896 parameter sets distributed across 25 topologies in Figure 2D). (B) Location of minimal kinetic filters in 3D space of first 3 principal components of 6 phenotypic metrics. See Figure S3 for singular values and composition of each principal component. Each sphere is centered at the mean principal component value observed over all kinetic filtering parameter sets of each minimal kinetic filtering architecture. Sphere size is proportional to radius capturing 60% of observed phenotypes. (C) Minimal kinetic filters cluster into 5 phenotypic groups that each share structural features. Archetypal topologies (right column) are the simplest topology in each phenotypic group.

Principal component analysis was performed on these phenotypic metrics for the set of minimal kinetic filtering topologies (**Fig S3**). **Figure 3B** presents each minimal topology as a sphere in principal component space with center at the mean principal component value across the topology’s kinetic filtering parameter sets and radius reflecting the amount of phenotypic variation within that topology.

The minimal kinetic filters fall into five functional clusters (**Fig 3C**). Principal component 1, consisting mainly of the long-term memory metric, divides the minimal kinetic filters into two groups, one with long-term memory and one without. The circuits with long-term memory are irreversible and bistable and are further divided by principal component 2 into two subtypes: ***positive feedback circuits with OR logic*** (**PFB**_OR_) and a class that we refer to as ***bistable inverters*** (**BI**).

Among the circuits without long-term memory, principal component 3 distinguishes between a class of kinetic filters that turn off slowly and gradually, requiring a long refractory period **-- *positive feedback circuits with AND logic*** (**PFB**_AND_) -- from kinetic filters that turn off quickly and steeply and are thus considered the most reversible. Principal component 2 further divides the reversible circuits into two subtypes, ***double inhibition*** (**DI**) circuits and ***coherent feed forward loops with AND logic*** (**CFFL**).

### Mechanisms of the five core kinetic filtering motifs

How does each of these five classes of core motifs achieve kinetic filtering behavior? Here we describe in detail the activation trajectories for each of these core classes, using ideal parameter sets that display kinetic filtering, and summarize their basic mechanisms. Parameter constraints observed for each of the five classes are summarized in **Fig S4**.

*Coherent feed forward loops (CFFL)* have been previously identified as being capable of kinetic filtering (Mangan and Alon, 2003). These topologies use a fast arm/slow arm mechanism for kinetic filtering, where the output node shows AND logic and is only activated if it receives simultaneous signals from both the fast arm (a measure for whether the input is still present) and the slow arm (a measure of whether input was also on some time ago). A representative coherent feed forward loop timecourse (**Fig 4A**) shows that output node C is only activated when both active A and active B are above a threshold concentration; thus, trigger time is largely determined by the time required to transmit signal through the slower arm of the network, or the difference between the slow and fast arms. Short inputs are filtered because they do not last long enough to allow both the long and short arm of the network to be simultaneously activating the terminal AND node (node C). CFFL circuits are reversible after removal of input, and turn off sharply. They can, however, display a moderate lag in initiating shutoff, which is dependent on how long the pool of active node A remains after removal of input.

**Figure 4.**
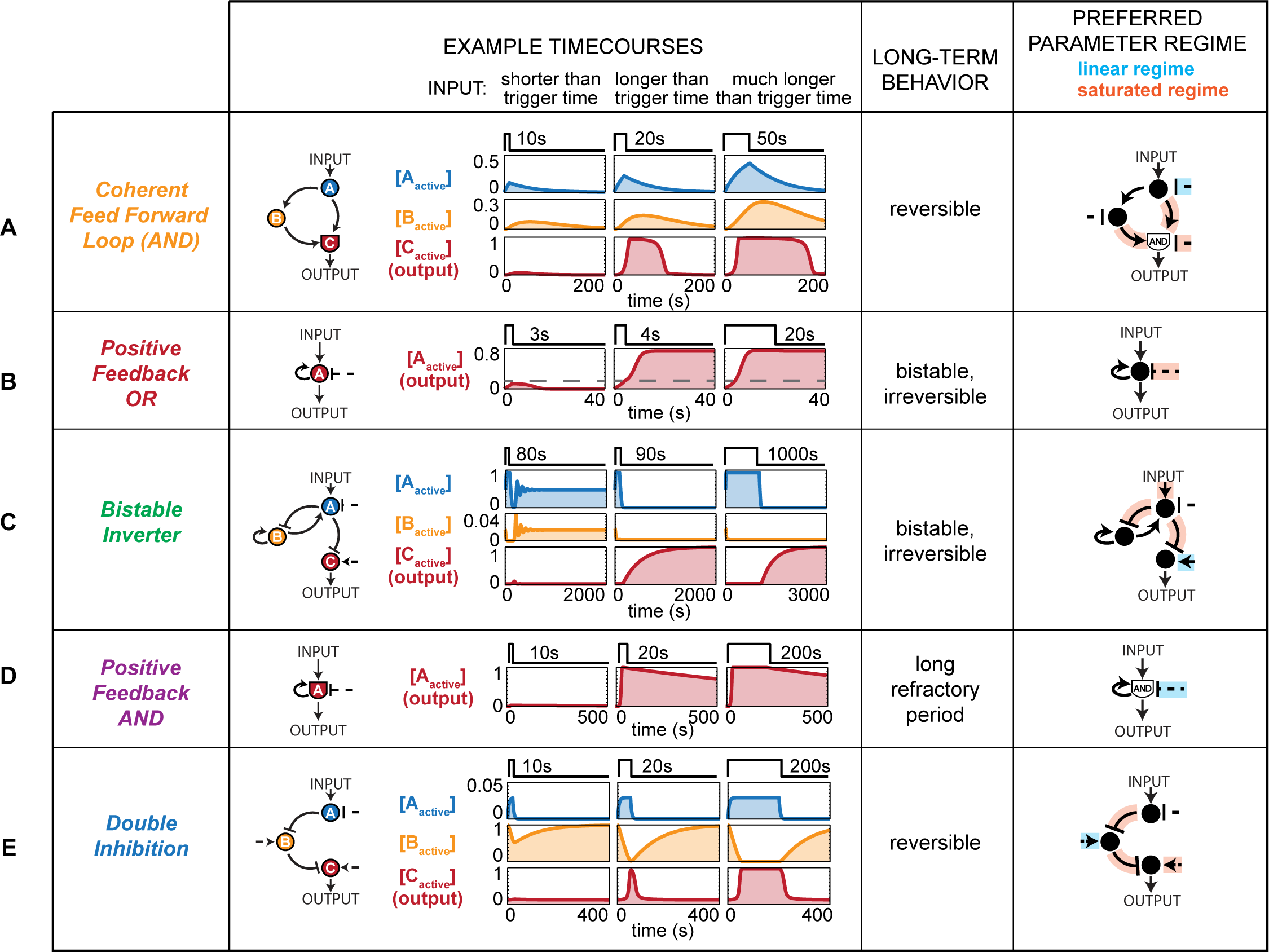
Representative timecourses and preferred parameter regimes of five classes of kinetic filters. Each circuit is shown responding to input shorter than trigger time, just longer than trigger time, and far longer than trigger time. Distributions of parameter values used to identify preferred regimes are shown in Figure S4. See Table S1 for parameter values used in example timecourses.

Bistable memory circuits, such as the *positive feedback OR circuit (PFB_OR_)*, can also achieve kinetic filtering. These circuits simply require a given time of input stimulation to pass the tipping point or separatrix, resulting in switching from the OFF state to the ON state (**Fig 4B**). The *PFB_OR_* motif requires only a single node to achieve a bistable circuit. Here activation of the OR node (node A) by input results in positive feedback stimulation to the node, which, once at sufficiently high levels to counter the basal inactivating activity, can self-sustain activation of the OR node even in the absence of input. Short inputs are filtered because they leave the level of activated node A below the threshold required for locking ON.

Another kinetic filtering motif that results in long-term memory is the *bistable inverter* (BI) class (**Fig 4C**). This highly unusual class of circuits exhibits behavior where the output only switches ON after input has been halted. To serve as a kinetic filter, the A and B nodes begin in activated states. The output node (node C) is thus initially off, because node A represses C. Upon input stimulation, the negative feedback relationship between B and A nodes initiates oscillations in the activity of both A and B, but if stimulation is long enough, then B becomes completely deactivated. In the absence of active B, when input is halted, the activity of node A falls to zero. Because node A was the only deactivator of the output node C, output will now turn on. In short, this type of kinetic filter requires a minimal input stimulation duration to “prime” the system and eliminate active B (**Fig S5**). This priming time defines the minimal trigger time of the kinetic filter. The priming period can be followed by a holding time of variable length as long as input is sustained, but immediately after the input is switched off, the output will turn ON irreversibly. This highly unusual motif has not been, to our knowledge, characterized in known examples of kinetic filters, but it is possible that this sort of two-phase switch may be useful for particular biological behaviors.

*Positive feedback loop AND (PFB_AND_)* circuits are another simple class of motifs capable of kinetic filtering (**Fig 4D**). In this case the integrating node (node A) must be simultaneously stimulated by input and positive feedback in order for the system to strongly activate output. Initial input can lead to low levels of activation of A, if the basal deactivators of A are weak (here an “AND” gate is not an absolute Boolean gate, but one in which the integration of two activating stimuli is multiplicative rather than additive). Nonetheless this buildup of initial amounts of activating A can be very slow and tuned by the strength of the basal opposing activity. If the input is on long enough to build up enough active A to trigger positive feedback, then A will turn on synergistically because of dual AND activation. In this case, the system does not formally display memory after input is removed, since it does turn off eventually. However, most parameters that lead to kinetic filtering also lead to an extremely slow inactivation of the system. Here the biggest difference between the PFB_AND_ and PFB_OR_ circuits is that the AND motif has a much stronger dependence on positive feedback in order to increase the level of active A. It is important to remember that our coarse grained search considered only two types of integrating node behaviors, and it is certainly possible that there could exist related positive feedback circuits in which the key integrating node has intermediate behaviors such as an OR gate where the weights of activation are much stronger for the positive feedback stimulation compared to the direct input stimulation. In this case, one would expect to obtain a kinetic filter that was like the PFB_AND_ motif in that it would require a long period of direct input stimulation to initiate build up of active A, but also like PFB_OR_ in that it could eventually lock on in a fully bistable manner.

The final major class of minimal kinetic filters is the *double inhibition (DI)* motif (**Fig 4E**). Networks in this class share a cascade with two successive inhibitory regulatory relationships. The central node (here B) starts with high activity and acts as an inhibitor of system output (node C). Input stimulation inhibits the inhibitor to switch on the system. DI motifs can act as kinetic filters when the central inhibitor node (here node B), acts as a buffer to absorb system input. Although input might immediately decrease node B activity, this does not register as a change in C node activity until after a longer duration of stimulation (trigger time) when the level of B has dropped very low,. Trigger time is determined by the overall rate of decrease in B node activity, which is dependent not only on input, but is also opposed by the basal activation of B; higher basal activation can yield longer trigger times. DI motifs are reversible: after removal of input, the system rapidly returns to its initial steady state. The DI motif can be arranged as a sequential element between input and output nodes as described above, but can also be found in a double inhibition feedback loops as long as a terminal element of the DI motif has a positive regulatory relationship with the output node.

### The most robust kinetic filtering networks are enriched for DI and PFB_AND_ motifs

To identify which of the five core motifs can give rise to the most robust kinetic filtering networks, we imposed a high-stringency robustness cutoff of 0.08 on the ~70,000 sampled topologies (**Fig 5A**). Remarkably, all 83 circuits in the high-robustness set contained DI and/or PFB_AND_ motifs but none of the other three core architectures (**Fig.5B**). Of the 83 highly robust kinetic filters, 25 have only PFB_AND_ motifs, 9 have only DI motifs, and 49 have both PFB_AND_ and DI motifs. The distributions of robustness of architectures containing each of the five core motifs (**Fig 5C**) show that circuits containing DI and PFB_AND_ motifs achieve much higher degrees of robust kinetic filtering than circuits containing CFFL and PFB_AND_ motifs, and circuits containing BI motifs are the least robust. The strong enrichment for both DI and PFB_AND_ motifs in the most robust kinetic filtering architectures prompted us to explore why these motifs may act as better kinetic filters than other related motifs.

**Figure 5.**
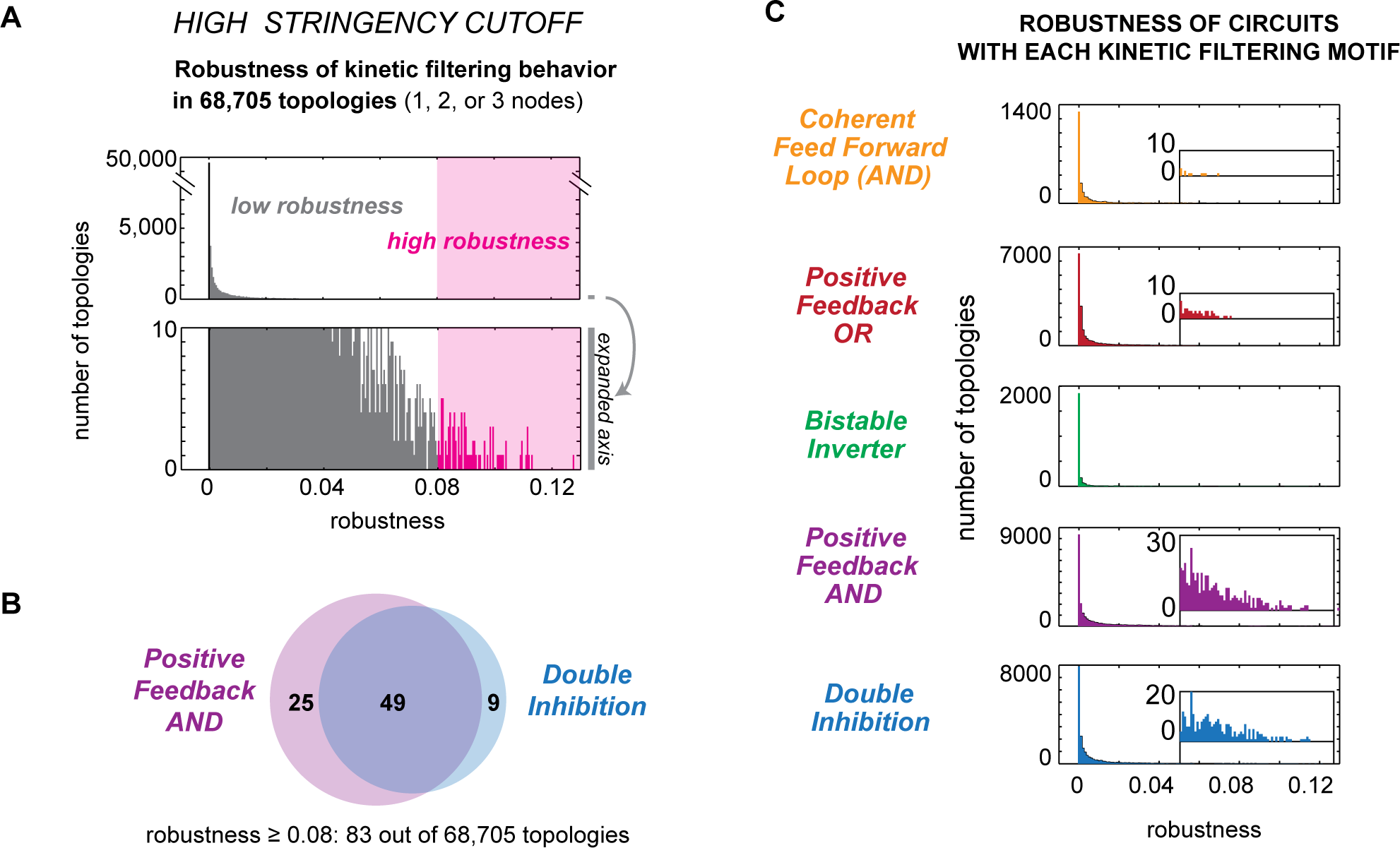
All highly robust kinetic filters include Positive Feedback AND and/or Double Inhibition motifs. (A) Distribution of robustness across all 68,705 enumerated topologies. Topologies with low robustness have robustness between 0.001 and 0.08; high robustness topologies have robustness ≥ 0.08. (B) Among the 83 topologies with high robustness, all contain either the positive feedback AND or the double inhibition motif, and most contain both. A circuit was considered to contain a kinetic filtering motif if it contained at least one of the minimal motifs in Figure 3C as a circuit substructure. (C) Distribution of robustness across all topologies containing a minimal kinetic filtering motif. Each topology was tested for containing each minimal kinetic filtering topology. A topology was considered to contain a double inhibition motif if it contained at least one double inhibition minimal kinetic filter, and analogously for each of the other kinetic filtering classes. All topologies that do not contain motifs in any of the 5 classes have robustness < 0.001.

### Why double inhibition cascades are better kinetic filters than double activation cascades

The more robust DI motif and the less robust CFFL motif are structurally analogous – the slow arm of the CFFL motif, which plays a major role in trigger time, is a double activation cascade, in contrast to the double inhibition motif. We examined the trigger time distributions of all DI parameter sets above our functional cutoffs, and found that they display trigger times ranging from 1s to >10,000s. Under the same range of sampled parameters, CFFL circuits are limited to trigger times under 100s (**Fig 6A**). When we examine the trigger times and temporal ultrasensitivity found with 10,000 random parameter sets imposed on archetypical DI or CFFL architectures, we find that DI circuits can occupy the quadrant with both high trigger time and high temporal ultrasensitivity (**Fig 6B**). In contrast, the CFFL circuits appear to have constraints that lead to a tradeoff between temporal ultrasensitivity and trigger time in the parameter sets that lead to kinetic filtering (**Fig S6**).

**Figure 6.**
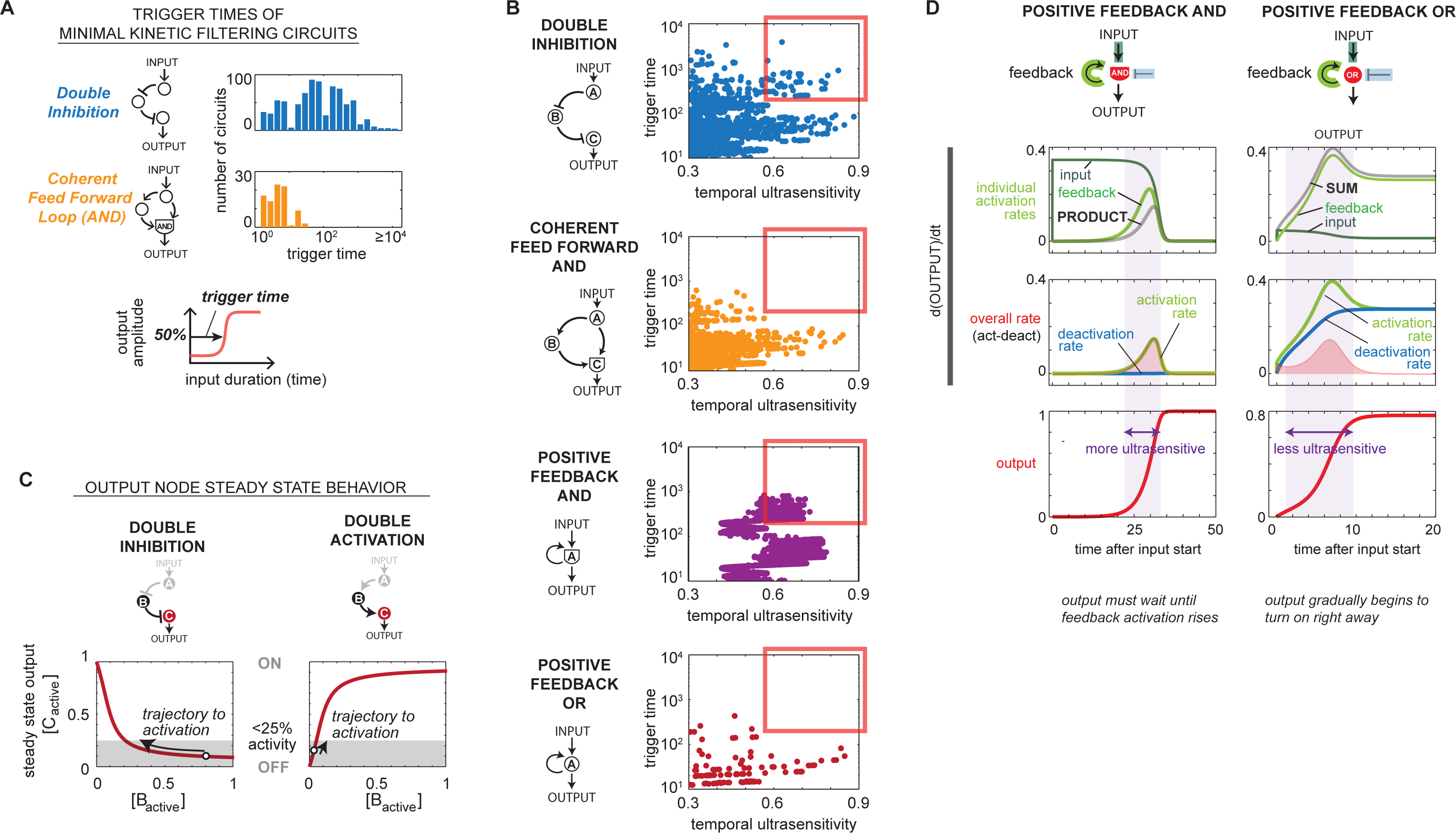
Turning off a deactivator more effectively buffers against partial activation by subthreshold length inputs. (A) Histogram of observed trigger times minimal kinetic filtering circuits. Parameter sets of minimal double inhibition topologies (#4, #21-24 in Figure 3C, total 798 parameter sets) and the coherent feed forward loop topology (#25 in Figure 3C, total 88 parameter sets) satisfying temporal ultrasensitivity score ≥ 0.5 and trigger time ≥ 1s were measured for trigger time. (B) Steady state output changes at a more gradual pace with changing regulator concentration in double inhibition circuits compared to double activation circuits. In both circuits, we solved for steady state output node concentration as a function of [B] with KmBC = 0.5, kcatBC = 1, Km basal activator/deactivator = 0.5, kcat basal activator/deactivator = 1, concentration of basal act./deact. = 0.1. (C) Positive feedback AND circuits are better kinetic filters than positive feedback OR circuits because they require output to remain low until feedback activation rises. Activation rate consists of activation due to input and activation due to feedback, which are multiplied in AND circuits and summed in OR circuits. Shaded region delineates the zone between 5% and 95% output activation. (D) An archetypal architecture for each kinetic filtering motif was sampled for 50,000 parameter sets over the same range as the sampling used in the enumerative search (kcat 0.1 to 10, Km 0.001 to 100, evenly in log space by Latin hypercube). Shown in each plot are the temporal ultrasensitivity score and trigger time for each parameter set of the archetypal topology that resulted in temporal ultrasensitivity score ≥ 0.3 and trigger time ≥ 10s (DI: 1328 parameter sets; CFFL: 1078 parameter sets; PFBAND: 2135 parameter sets; PFBOR: 115 parameter sets).

These differences between the behavior of DI and CFFL kinetic filters likely result from intrinsic differences between turning on an output by activating an activator versus inhibiting an inhibitor. To illustrate this point, we directly contrast a double inhibition circuit with a double activation circuit modeled with identical parameters — a double activation circuit is simply a coherent feed forward loop with the short arm removed. In **Fig 6C** we solve for the steady state concentration of active output node (node C) as a function of the fraction of active regulator node (node B). In both cases, output is initially low prior to input and increases the longer input is applied. For double inhibition circuits, the shape of the steady state output curve dictates that output remains low for a wide range of node B concentration; only after the amount of active B has been driven below a threshold does the fraction of active output (node C) begin to rapidly increase. Thus for double inhibition circuits, trigger time can be tuned to be very long, with little partial activation before reaching the trigger time. In contrast, for double activation circuits, output begins to partially increase substantially for even a small amount of initial increase in node B. Trigger time is thus limited by the shape of the output activation curve being steepest at low concentrations of node B, the early phase of the stimulation trajectory.

This simple observation that double inhibition cascades will have intrinsically distinct temporal activation properties from double activation cascades is related to earlier work from Savageau on the noise resistance of different regulatory schemes (Savageau, 1977). Systems that switch on by double inhibition cascade are more noise-resistant than systems that switch on by activation cascade because of the intrinsic difference in which regimes of the dose-response curve are steep or shallow.

### Why Positive Feedback AND motifs are better kinetic filters than Positive Feedback OR motifs

We then compared why the PFB_AND_ motif is a more robust kinetic filter than the closely related PFB_OR_ motif. Taking the archetypical kinetic filter of each architecture and carefully dissecting the trajectory of output activation, we observe that for the PFB_AND_ motif, no significant amount of output activation occurs until the feedback activation loop has been significantly triggered, as would be expected for the AND integration of the central node (**Fig 6D**). Feedback activation can trigger only very late in the trajectory because activation requires accumulation of the activated node, which can only occur through leaky activation induced by direct input activation.

In contrast, the PFB_OR_ circuit is able to immediately show significant and gradual activation of the central node, because the OR integration allows significant activation by the direct input even in the absence of positive feedback. Thus the PFB_OR_ circuit activation trajectory is inherently less steep than the equivalent PFB_AND_ circuit.

## DISCUSSION

### General classes of signaling networks capable of kinetic filtering and their potential functional roles

Here we have used coarse-grained network enumeration to identify classes of biochemical networks that can achieve kinetic filtering – the ability of a system to respond only after input stimulation has sustained for a given threshold duration. Networks that execute kinetic filtering can play a central role in filtering transient noise, interpreting complex dynamic inputs, and controlling the timing of a sequence of events. Given the growing appreciation of the importance of dynamics in cellular information processing, it will be important to understand the mechanisms that can be used for kinetic filtering and recognize and classify the networks that are found in cells (Purvis and Lahav, 2013). The design principles of such kinetic filters may also allow us to design cellular circuits with precision temporal control (Lim et al., 2013).

Kinetic filters must be able to absorb and dissipate input pulses that are shorter than a threshold triggering time, thus suppressing the resulting output. Here we focus on two behavioral parameters of the network: the *trigger time* -- the threshold stimulus duration time required to achieve half-maximal output, and *temporal ultrasensitivity* – the steepness of system activation as a function of input duration. Ideal kinetic filters can be considered to have both high temporal ultrasensitivity and high trigger times.

Our metric of temporal ultrasensitivity is analogous to the cooperativity index used to describe concentration-based ultrasensitivity. While well-established, this metric is sensitive to right translations: a temporal dose response of identical steepness will have a poorer score if trigger time is higher. Our analysis has focused on general properties of broad classes of circuits, but future work on kinetic filters may benefit from exploring other metrics capable of distinguishing subtle behaviors such as thresholding and switching (Gunawardena, 2005).

When we perform an exhaustive search for networks capable of kinetic filtering, we identify five classes of architectures. Some classes (PFB_OR_, CFFL) require tradeoffs between high temporal ultrasensitivity and long trigger time, while others (PFB_AND_, DI) allow simultaneous optimization of trigger time and temporal ultrasensitivity. These findings suggest different potential functional roles for these different classes of circuits (**Fig 7A**). The CFFL can effectively filter against activation by relatively transient noise, since this would only require optimization of a high temporal ultrasensitivity without a long trigger time. DI and PFB_AND_ circuits may be better as kinetic filters that also incorporate a longer timer or delay function, since they can exhibit long trigger time without sacrificing temporal ultrasensitivity. Finally, PFB_AND_ circuits could also be used in cases where memory or a long turn-off lag is needed, while CFFL and DI would be more suited to cases where rapid shut-off is optimal.

**Figure 7.**
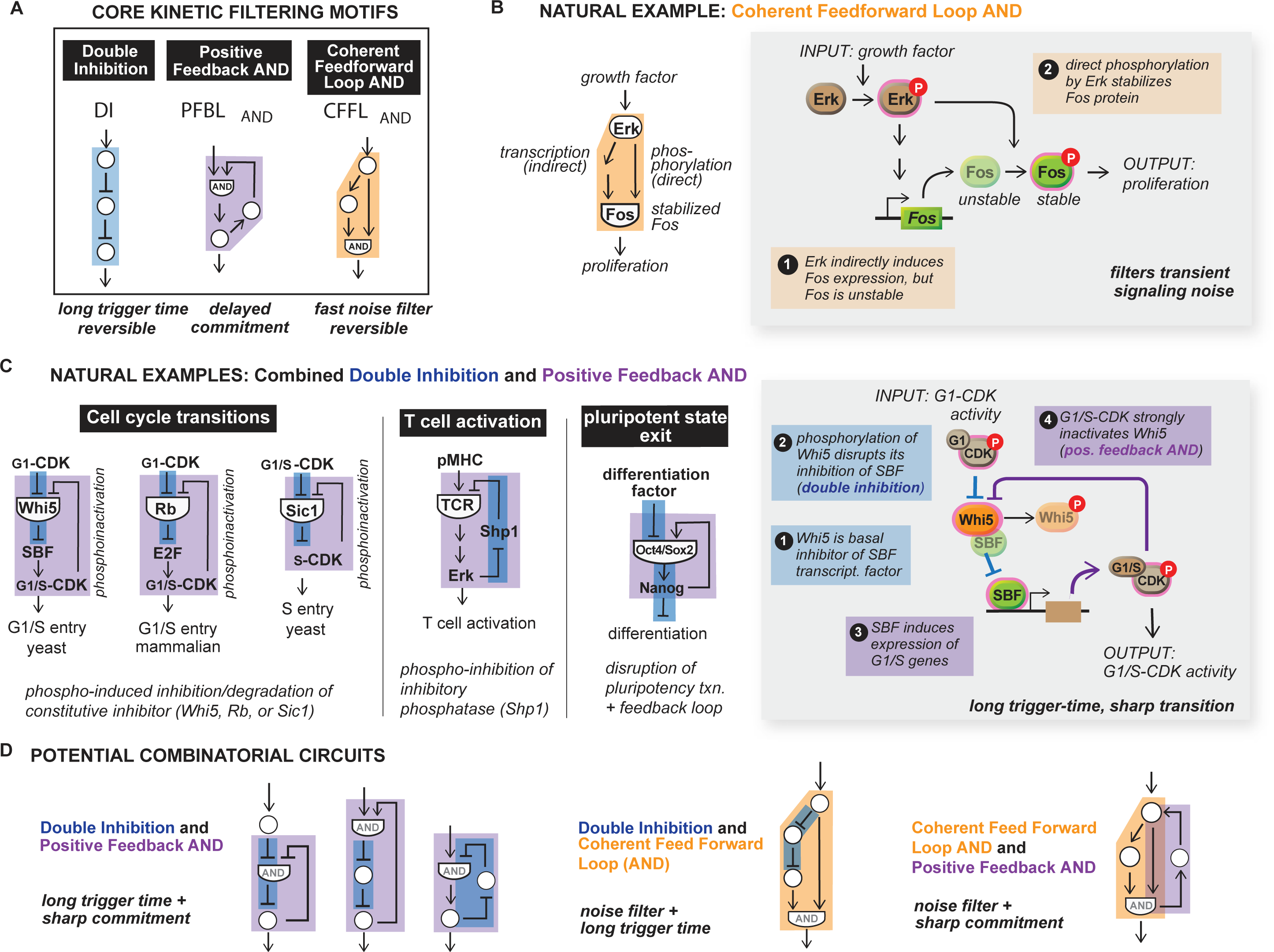
Natural examples of kinetic filters feature both core and combinatorlal kinetic filtering motifs. (A) Double inhibition, positive feedback AND and coherent feed forward loops form the core set of kinetic filtering motifs. (B) Growth factor response in PC-12 cells is governed by a kinetic filter with coherent feed forward AND architecture implemented through transcription and phosphorylation. (C) Cell cycle transitions are controlled by double inhibition and positive feedback AND architectures in both mammalian and yeast cells. T cell activation and pluripotent state exit in embryonic stem cells use a double inhibition / positive feedback AND circuit for kinetic filtering of short input signals. (D) Core kinetic filtering motifs can be combined to yield hybrid phenotypes.

### CFFL motifs in natural kinetic filtering circuits: Erk activation of cFos

This analysis predicts that one should be able to find these circuit types in natural kinetic filtering systems, although some might be preferred for a given functional context. There are several signaling systems that are known to display kinetic filtering. Here we examine these natural evolved systems and compare them with the motifs identified in this theoretical study.

One classical example of kinetic filtering is the activation of the cFos protein only in response to sustained activation of the mitogen activated protein kinase (MAPK) Erk (**Fig 7B**). In this case, activation of cFos occurs through a CFFL network (Murphy et al., 2002). Erk activation of transcription factors leads to increased transcription of the cFos gene. The cFos protein is, however, rapidly degraded, so it does not accumulate. Erk also directly phosphorylates the cFos protein, resulting in cFos stabilization. Here Erk-mediated transcription of cFos serves as the slow branch of the CFFL network, while direct Erk phosphorylation of cFos serves as the fast branch. cFos accumulation acts as an AND gate, since both cFos transcription and phosphorylation are required.

It is hard to know exactly why the somewhat less robust CFFL architecture is used in this case. One possibility is that in some cases (e.g. EGF stimulation), Erk activation occurs in pulses, where the frequency of pulses can convey information (Albeck et al., 2013). For a downstream output to effectively integrate multiple Erk pulses only when they are relatively close together would require a system that neither shuts off immediately nor has an extremely long lag time. The CFFL may be ideal for this kind of frequency-encoded information, since the DI networks shut off extremely fast, and the PFB_AND_ networks show very long turn-off lag.

### Convergence on combined DI and PFB_AND_ motifs in cell cycle transition control networks

Several natural kinetic filters seem to have converged upon a similar combination of both the DI and PFB_AND_ motifs, the two most robust kinetic filtering architectures identified in this work. Recent studies have observed that several phase transitions in the cell cycle utilize convergent regulatory networks (**Fig 7C**) (Bertoli et al., 2013; Skotheim et al., 2008; Yang et al., 2013). In each of these cases, the networks contain integrated DI and PFB_AND_ motifs. In key cell cycle transitions, the cell starts with high activity of the cyclin dependent kinase (CDK) in complex with an initial phase. The cell must then sharply transition to the next phase, associated with a sharp increase in the next phase cyclin-CDK complex. These combination DI and PFB_AND_ networks appear to be optimal to drive this transition in a temporally sharp and decisive manner.

All of these circuits, even though they operate at different stages of the cell cycle, or in different organisms, have a central inhibitor that initially prevents output, i.e. next phase cyclin-CDK activity. For yeast entry into START (the G1/S transition), the inhibitor is the protein Whi5, which binds to and inhibits the transcription factor SBF. The transition initiates with a double inhibition cascade: when Whi5 is phosphorylated by the initial phase G1/Cdk enzyme, it initiates release from SBF, which in turn allows SBF to initiate expression of G1/S phase genes, including the G1/S cyclins. This results in increase in G1/S Cdk enzyme, the next phase cyclin/CDK complex, which then acts in a strong positive feedback manner to even more strongly phosphorylate and inactivate Whi5. Here, because Whi5 requires priming phosphorylation by the G1-CDK complex, but is more efficiently phosphorylated by the G1/S Cdk enzyme, it approximates an AND gate. Overall, this system shows a very sharp temporal transition after a long delay, followed by a strong commitment to the next phase, a combination of behaviors that the DI/PFB_AND_ hybrid network should be ideal for. Here the central inhibitory node (Whi5 = node B) is not an enzyme, as in the case of the network models used in our coarsegrained search, but is instead a stoichiometric inhibitor. This particular molecular manifestation of the network is expected to show temporal ultrasensitivity as long as the binding of Whi5 to SBF is sufficiently tight (**Fig S7**).

Strikingly, the identical hybrid network is observed in other cell cycle transitions. In mammalian G1/S entry, the protein Rb serves as the central inhibitory node analogous to Whi5, even though it is evolutionarily unrelated (Bertoli et al., 2013). Rb is an inhibitor of a transcription complex, and Rb’s function is in turn initially inhibited by G1-CDK mediated phosphorylation. Rb’s inhibition leads to expression of the G1/S cyclins, leading to positive feedback when the G1/S CDK enzyme strongly phosphorylates Rb.

In the case of the yeast S phase entry, the protein Sic1 serves as a central inhibitory node. Sic1 is a direct inhibitor of the S-Cdk complex, but is initially phosphorylated by the earlier stage G1/S-CDK enzyme. In a DI motif, phosphorylation of Sic1 leads to its degradation, initiating activation of the next phase G1-CDK enzyme. This leads to positive feedback, since the G1-CDK enzyme more strongly phosphorylates Sic1, leading to its even more rapid degradation.

### Combined DI and PFB_AND_ network in T cell activation and stem cell exit from pluripotency

Committed activation of T cells upon antigenic peptide-MHC engagement is thought to involve kinetic filtering (Davis et al., 1998). Activation is only observed with peptide-MHC complexes with sufficiently long engagement times. One of the key proteins thought to play a role in this kinetic filtering is the negatively regulatory phosphatase Shp1 (Altan-Bonnet and Germain, 2005; Feinerman et al., 2008). Examination of the Shp1 network reveals a combined DI and PFB_AND_ circuit (**Fig 7C**). Shp1 acts as an inhibitor that removes activating phosphorylation on the T cell receptor (TCR) and some of its downstream effectors. Activation of the TCR leads to its own phosphorylation, and in subsequent steps, activation of the downstream MAPK Erk. Active Erk can in turn phosphorylate Shp1 in a manner that is thought to lead to its dissociation from the TCR complex. Thus this network contains a DI cascade integrated with a positive feedback loop, with a related but slightly distinct configuration from the combined DI and PFB_AND_ networks observed in the cell cycle transitions discussed above.

Similarly, the differentiation of pluripotent embryonic stem cells only to sustained but not transient differentiation signals (Sokolik et al., 2015). The circuit that induces differentiation (or exit from the pluripotent state) has a combined DI and PFB_AND_ motif. The Oct4-Sox2-Nanog maintains the pluripotent state through an autoregulatory positive feedback loop, thus acting as a repressor of differentiation. Differentiation factors induce the switch by disrupting the Oct4-Sox2-Nanog complex through competitive binding, leading to Nanog degradation and thereby relief of the repression of differentiation (double inhibition) (**Fig 7C**).

### Other potential combinatorial kinetic filtering networks

Some of the best-characterized cellular systems that display kinetic filtering contain combinations of the ideal core motifs identified in our analysis. We predict several other possible combinatorial motif circuits to have useful combinations of behaviors. For example, a DI and CFFL combination circuit (**Fig 7D**) is expected to yield both a long trigger time and intermediate off-kinetics. Such a circuit could be used to integrate multiple wide pulses of input. Combining CFFL and PFB_AND_ could lead to efficient transient noise filtration combined with a committed transition. These networks built of multiple combinations of the core kinetic filtering motifs require more than three nodes, and thus would not have been identified from our enumeration of 3-node networks.

It is likely that the core motifs identified here could be combined together, both sequentially or in an interlinked manner to build even more effective kinetic filters with longer trigger times, or in a way that can overcome particular functional tradeoffs of the individual simpler motifs. We have previously found that a similar combinatorial use of minimal motifs leads to more robust cell polarization circuits (Chau et al., 2012). Multiple kinetic filters could be linked together in higher order sequences of events that control processes like the cell cycle or trafficking that require distinct steps to occur in a defined order. It is also possible that these minimal motifs could be combined with oscillatory networks to yield timer systems that combine pulsatile clock-like mechanisms with kinetic filtering (Suel et al., 2007).

### Evolutionary choice of signaling enzyme regulatory mechanisms may be linked to dynamic response behaviors

The diverse dynamic behaviors examined here may also explain why particular molecular mechanisms of regulation are chosen for different signaling enzymes. One of the most prevalent molecular mechanisms to gate signaling enzyme activity is regulation via an inhibitory domain that can act in *trans* or in *cis* (autoinhibition). Activation can thus occur via double inhibition or relief of autoinhibition. Although these mechanisms of enzyme regulation are very similar at a molecular level, this study suggests that when incorporated into circuits, the two molecular switches will have very different dynamical properties. At a network level, circuits with core nodes that rely on regulation by relief of autoinhibition will behave like conventional activation cascades, which can switch on faster but are less likely to have sharp temporal ultrasensitivity. In contrast, systems with an unlinked inhibitor could achieve much sharper temporal ultrasensitivity. It will be interesting to explore whether known signaling systems that use *trans* inhibition (e.g. protein kinase A, which is regulated by an inhibitory subunit that dissociates upon cAMP binding) are associated with robust time delays, while those that use *c* -inhibition (e.g. Src kinases, which are regulated by intramolecular autoinhibitory domain interactions) are associated with faster, more immediate processes.

## CONCLUSIONS

The motifs and understanding that emerge from this enumerative circuit analysis provide a useful roadmap for more deeply and predictively understanding how cells interpret dynamic information. These mechanisms can help understand why particular network perturbations that might disrupt timing control could contribute to diseases such as cancer. The motifs that emerge also provide a catalog by which to define key dynamical control elements within complex cellular networks mapped by proteomic and genomic methods.

## METHODS

### Simulation of biochemical circuits

Reactions were modeled with total quasi-steady-state Michaelis-Menten kinetics (Ciliberto et al., 2007; Gomez-Uribe et al., 2007; Tzafriri, 2003). Nodes were converted between active and inactive states according to network linkages, where positive regulations catalyzed activations and negative regulations catalyzed deactivations (Fig S1). The total concentration of each node was held constant at 1. For nodes operating under OR logic, Michaelis-Menten expressions for incoming links were added. For nodes operating under AND logic, Michaelis-Menten expressions for incoming links of the same sign were multiplied, and expressions for incoming links of opposite signs were added. Each circuit was numerically integrated with a fifth-order embedded Runge-Kutta formula (Press et al., 2002). Active concentrations of each node were initialized to 0.1 and allowed to come to steady state before the application of input.

To enumerate circuit topologies, we allowed each link to be positive, negative, or absent. We discarded topologies where the input signal did not reach the output node. Circuits with regulations on a non-input, non-output node that did not in turn regulate another node were also discarded. For AND logic topologies, we discarded all circuits where the node with AND logic did not have two regulatory links of the same sign, counting input as a positive regulation.

In addition to the regulations between nodes A, B, and C, a circuit had additional constitutive activators and deactivators as needed such that no node had only activators or only deactivators (Fig S1). Constitutive activators and deactivators had constant concentration of 0.1. Up to 26 parameters were sampled for each circuit: k_cat_ and K_m_ for each of the nine possible circuit links, three possible constitutive activators and deactivators, and the input link. Node concentration was held constant at 1.0 and not sampled. All parameter samplings used the Latin hypercube method (Iman et al., 1980) with range 0.1 to 10 for k_cat_ and 0.001 to 100 for K_m_; this range is roughly physiological with units of seconds and μM. 10,000 parameter sets were sampled for the enumerative search and 100,000 for determining parameter regime restrictions.

### Quantification of circuit performance

Input pulses of duration 0.25, 0.5, 1, 2, 3, 4, 5, 6, 7, 8, 9, 10, 20, 30, 40, 50, 60, 70, 80, 90, 100, 200, 300, 400, 500, 600, 800, 1000, 2000, 3000, 5000, 6000, 8000, 10000, 20000, and 50000 seconds were applied separately to each parameter set of each topology. Input amplitude was always 0.1. Maximum output amplitude was measured over the period covering both the duration of the input pulse as well as a post-pulse recovery period lasting until the system came to steady state. Circuits that failed to reach steady state within 86,400 simulation seconds were removed from consideration. Inverting circuits whose output decreased with application of input were also discarded.

Temporal ultrasensitivity was quantified by plotting the circuit’s maximum output amplitude for each duration of input and measuring the temporal ultrasensitivity score (TU score) of the resulting curve (Fig 1B). TU score was defined as the ratio of input duration yielding 10% of maximum response to input duration yielding 90% of maximum response, analogous to cooperativity score in classical dose response curves (Goldbeter and Koshland, 1981). The 10% and 90% input durations were determined by interpolating a linear fit between the simulated input durations bracketing the 10% and 90% response amplitudes respectively. The response threshold T was determined by linear fit between input durations bracketing 50% output amplitude.

Maximal response (Rmax) and difference between maximal and minimal response (ΔR) were also measured for each circuit. Circuits with Rmax < 0.001 or ΔR/Rmax < 0.5 were considered to have insufficient response amplitude and were not considered to be kinetic filters.

## AUTHOR CONTRIBUTIONS

JG and WAL conceived the project. JG designed and carried out the analysis. JG and WAL wrote the manuscript.

## ACKNOWLEDGEMENTS

The authors thank Chao Tang, Hana El-Samad, Wallace Marshall, Wenzhe Ma, Angi Chau, Matt Thomson, Russell Gordley, and Gabriel Rocklin for helpful discussions.

## SUPPLEMENTAL INFORMATION

**Figure S1.**
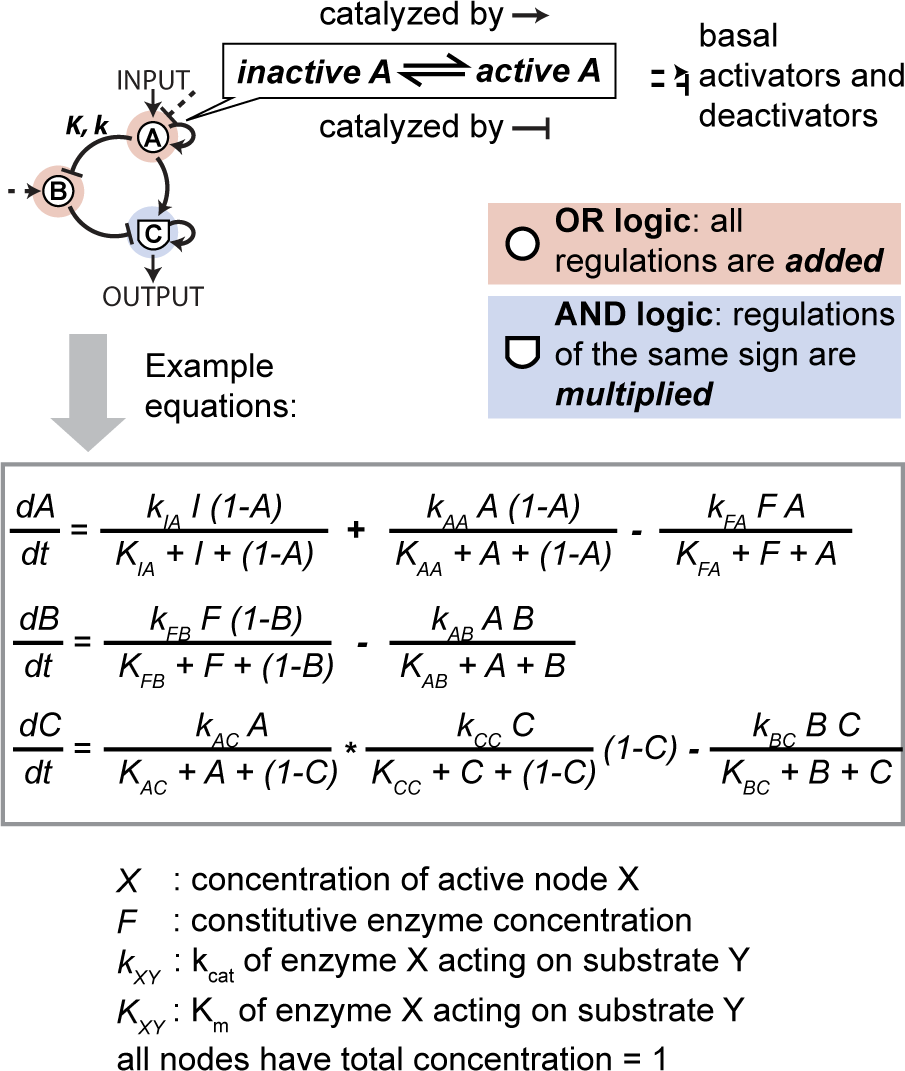
Modeling enzymatic circuits with OR and AND logic. Reactions were modeled with total quasi-steady-state Michaelis-Menten kinetics (tQSS-MM). Nodes were converted between active and inactive states according to network linkages, where positive regulations catalyze activations and negative regulations catalyze deactivations. The total concentration of each node was held constant at 1, and only the active fraction of each node could catalyze other reactions. For nodes operating under OR logic, tQSS-MM expressions for incoming links were added. For nodes operating under AND logic, tQSS-MM expressions for incoming links of the same sign were multiplied, and expressions for incoming links of opposite signs are added. Basal activators and deactivators were added as needed such that no node had only activators or only deactivators. Basal activators and deactivators had constant concentration of 0.1.

**Figure S2.**
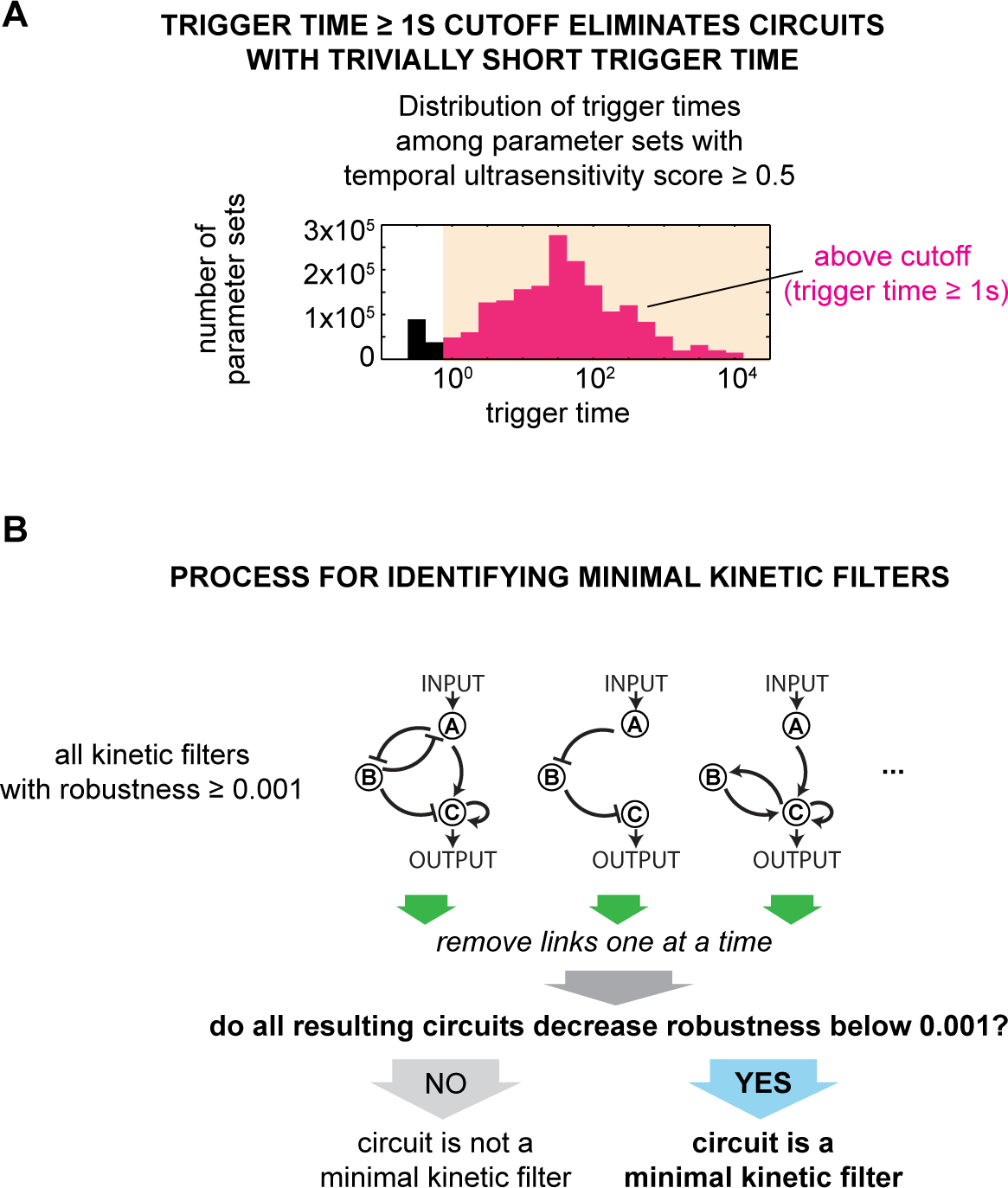
Characterizing kinetic filters. **A.** Distribution of trigger times among all parameter sets (from 68,705 topologies) with temporal ultrasensitivity score ≥ 0.5. Parameter sets with trigger time < 1s were discarded to avoid circuits with trivially short trigger time. **B.** Minimal kinetic filters were identified by removing each link from the set of topologies with robustness ≥ 0.001 and testing whether link removal resulted in a circuit with robustness < 0.001. Minimal kinetic filters are those where removal of any single or combination of links results in decreasing robustness below 0.001.

**Figure S3.**
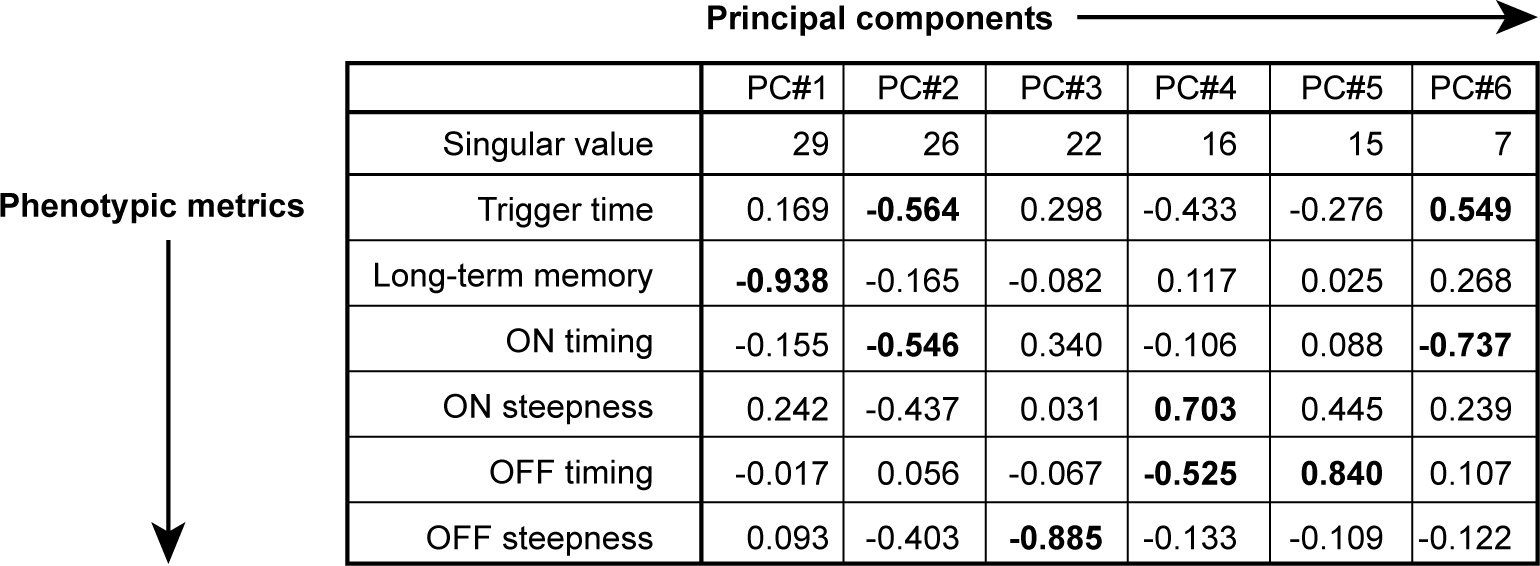
Composition and singular values of six principal components of phenotypic space. The square of the singular value reflects the amount of variation in phenotype explained by each principal component. The composition of each principal component is given in terms of relative weight of each phenotypic metric (Figure 3A). Primary contributors to each principal component (magnitude of weight > 0.5) are bolded. Principal components were identified with the SciPy linalg.svd package over 2,896 measurements of the 6 phenotypic metrics.

**Figure S4.**
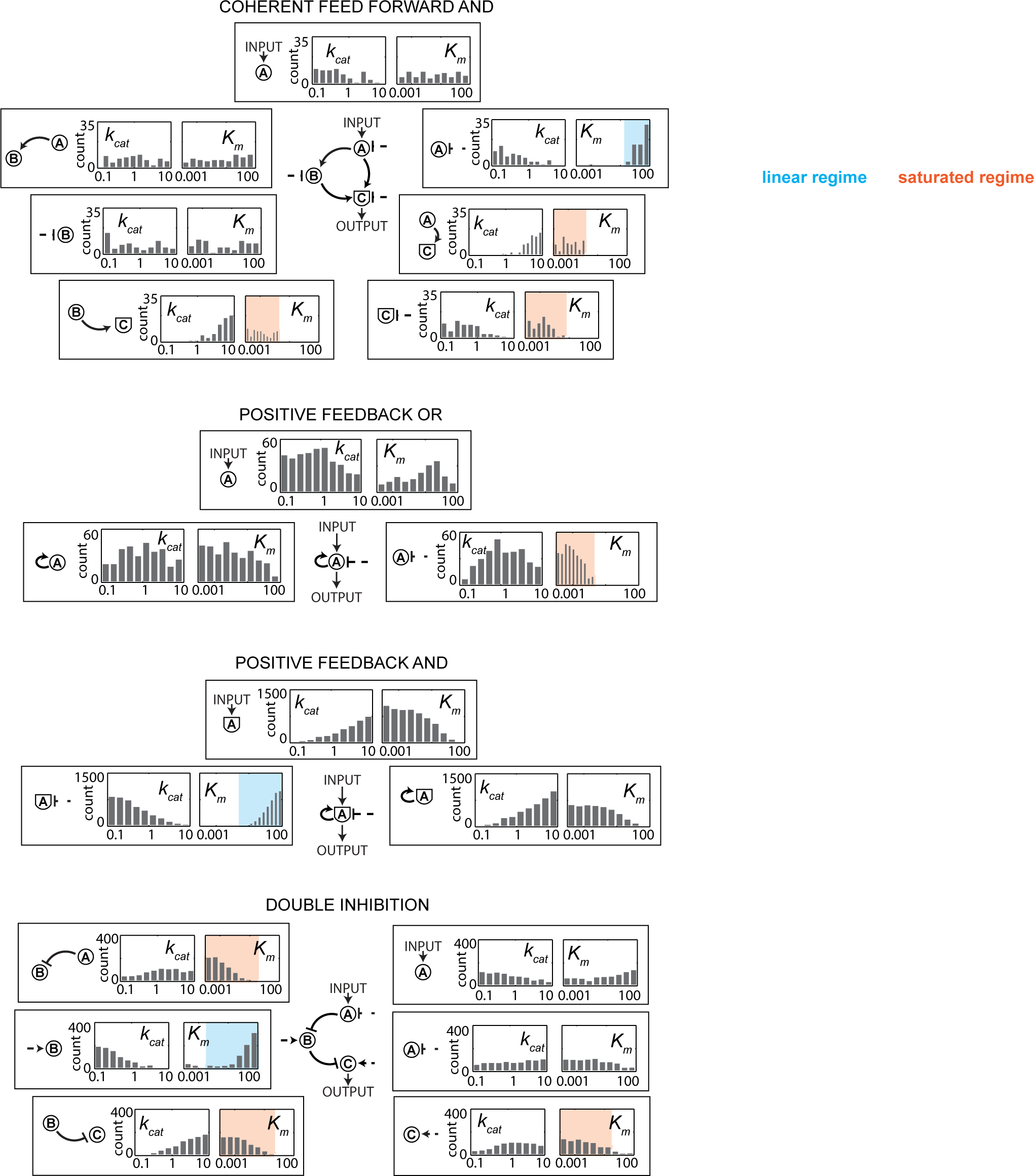
Preferred parameter regimes of minimal kinetic filtering circuits. An archetypal architecture for each kinetic filtering motif was sampled for 100,000 parameter sets over the same range as the sampling used in the enumerative search (kcat 0.1 to 10, Km 0.001 to 100, evenly in log space by Latin hypercube). Shown in each plot are the temporal ultrasensitivity score and trigger time for each parameter set of the archetypal topology that resulted in temporal ultrasensitivity score ≥ 0.5 and trigger time ≥ 1s (Coherent feed forward loop: 518 parameter sets; Positive feedback OR: 323 parameter sets; Positive feedback AND: 4391 parameter sets; Double inhibition: 907 parameter sets).

**Figure S5.**
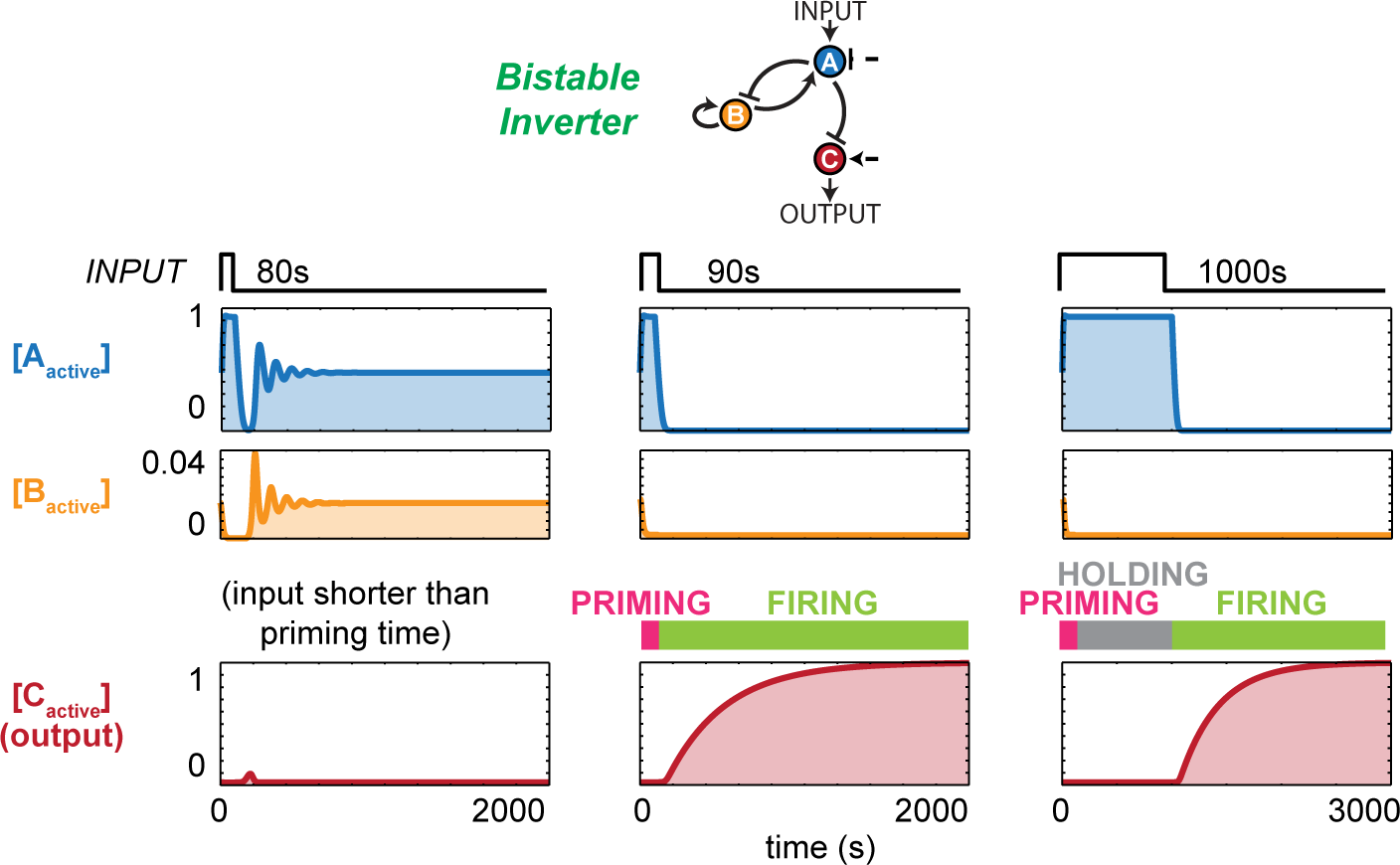
Bistable inverters respond by priming, holding, and firing. Upon input, bistable inverters begin a priming process during which node B begins to be inactivated. If input duration is shorter than the length of time needed for priming, i.e. the length of time needed to completely deactivate node B, the bistable inverter circuit does not fire. If input duration is longer than priming time, the circuit finishes priming and enters a holding phase during which output remains low. Once input has turned off, the circuit exits the holding phase and fires.

**Figure S6.**
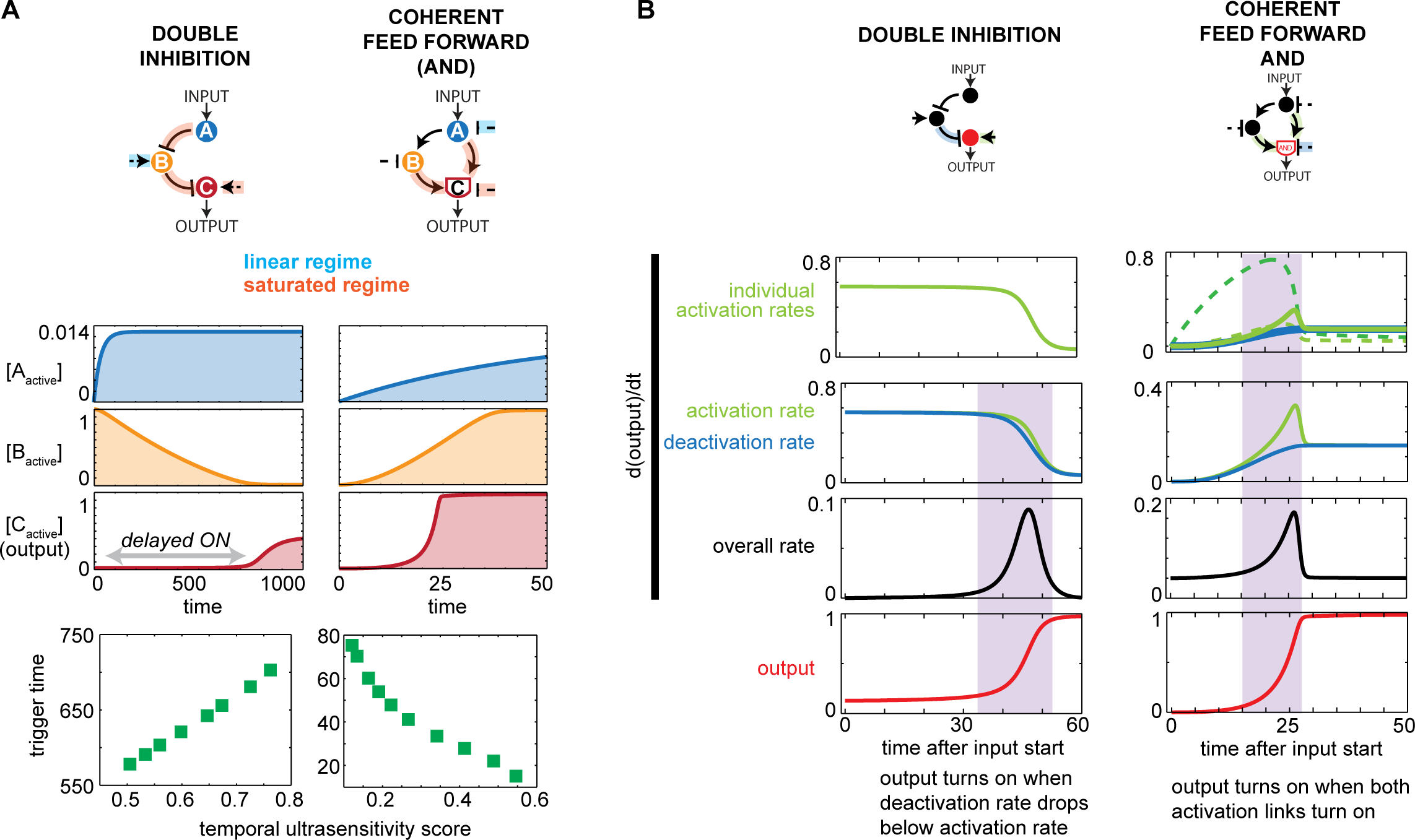
Double inhibition and coherent feed forward kinetic filter mechanisms. **A.** Trigger time and temporal ultrasensitivity score can be simultaneously optimized in DI but not CFFL circuits. For both circuits, a parameter set satisfying temporal ultrasensitivity score ≥ 0.5 and trigger time ≥ 1s was used to plot timecourses. For the DI circuit, the concentration of the basal activator of the output node C, usually held at 0.1, is tuned from 0.1 to 0.8, and trigger time and temporal ultrasensitivity scores of each resulting circuit are measured and plotted against each other. For the CFFL circuit, concentration of the basal deactivator of output node C, usually held at 0.1, is tuned from 0.1 to 1.0 and trigger times and temporal ultrasensitivity scores of the resulting circuits are plotted against each other. **B.** Activation and deactivation rates for DI and CFFL circuits (same parameter sets as in Figure 4E and 4A respectively). For the CFFL, activation terms of each node activating the output node are shown separately (tall dotted activation: fast arm; low dotted activation: slow arm); their product constitutes the overall activation term for output node C. Shaded region delineates the zone between 5% and 95% output activation.

**Figure S7.**
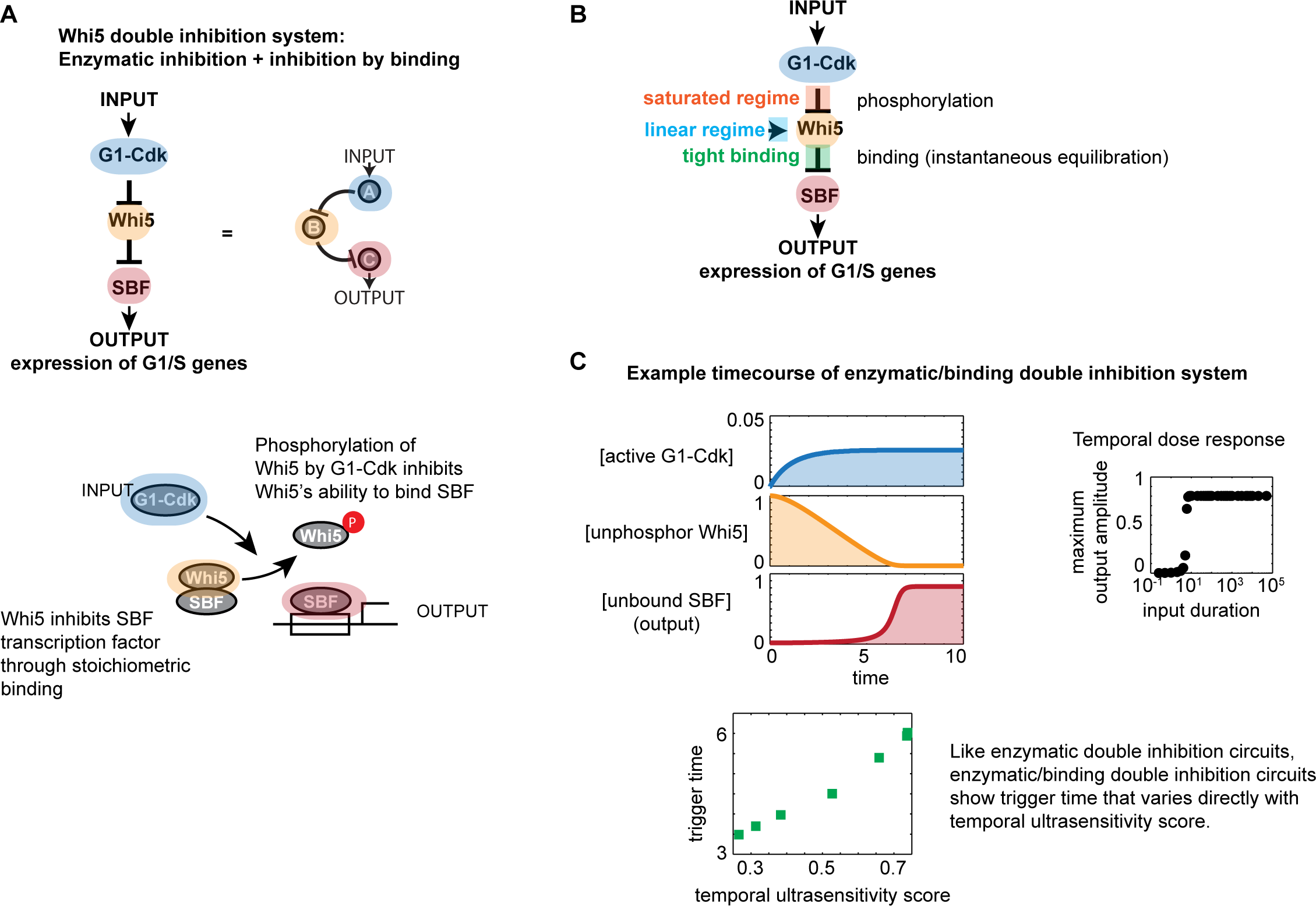
Double inhibition circuits using an enzymatic/binding mechanism can still function as kinetic filters. **A.** The yeast G1/S transition is a double inhibition circuit with the first inhibition enzymatically regulated and the second inhibition regulated by binding interactions. **B.** The Whi5 double inhibition system can be a kinetic filter. A double inhibition system was set up with enzymatic regulation of input activating G1-Cdk, G1-Cdk inactivating Whi5, and constitutive activator of Whi5. Active Whi5 equilibrates instantly with unbound SBF to form Whi5-SBF complex. Output measures the amount of unbound SBF. Parameters were sampled evenly in log space using Latin hypercube over the ranges 0.1 to 10 for k_cat_s, 0.001 to 100 for K_m_s, and 0.001 to 100 for K_d_ of Whi5-SBF binding. Parameter restrictions required for kinetic filtering behavior are shown as saturated regime for Whi5 phosphorylation, linear regime for Whi5 dephosphorylation, and tight binding for the Whi5-SBF interaction. **C.** A representative timecourse is shown here with its temporal dose response curve. Like fully enzymatic double inhibition kinetic filters, the enzymatic/binding double inhibition circuit can also be simultaneously optimized for long trigger time and steep temporal dose response.

**Table S1:** Parameters used for example timecourses in Figure 4.

| Circuit | Link | $k_{cat}$ | $K_m$ |
| --- | --- | --- | --- |
| Coherent feed forward loop (AND) | Input | 1.67109 | 9.98849 |
|  | AB | 0.196426 | 4.98884 |
|  | AC | 4.75554 | 0.149796 |
|  | BC | 2.90402 | 0.0959401 |
|  | A deactivator | 0.180053 | 0.724436 |
|  | B deactivator | 1.05779 | 4.43098 |
|  | C deactivator | 1.61287 | 0.0110917 |
| Positive feedback OR | Input | 2.09411 | 3.37287 |
|  | AA | 1.5531 | 0.06026 |
|  | A deactivator | 3.24638 | 0.0415 |
| Bistable inverter | Input | 0.565458 | 0.00248599 |
|  | AB | 3.85301 | 15.1705 |
|  | AC | 0.125314 | 0.00300608 |
|  | BA | 1.28529 | 0.0295801 |
|  | BB | 0.153957 | 0.296483 |
|  | A deactivator | 0.298813 | 0.168461 |
|  | C activator | 2.47172 | 86.3973 |
| Positive feedback AND | Input | 4.33112 | 0.14962 |
|  | AA | 0.94493 | 0.06173 |
|  | A deactivator | 0.10195 | 12.7057 |
| Double inhibition | Input | 1.71791 | 25.704 |
|  | AB | 1.03134 | 0.00245471 |
|  | BC | 4.96821 | 0.00389493 |
|  | A deactivator | 0.493856 | 0.0989692 |
|  | B activator | 0.29964 | 2.43781 |
|  | C activator | 7.67361 | 0.21208 |

